# Polyamines shift expression of macrophage L-arginine metabolism related-genes during *Leishmania amazonensis* infection

**DOI:** 10.1101/2022.05.27.493766

**Authors:** Jonathan Miguel Zanatta, Stephanie Maia Acuña, Camilla de Almeida Bento, Beatriz Simonsen Stolf, Sandra Marcia Muxel

## Abstract

Polyamines are molecules involved in macrophage activation and polarization in response to pathogens. The interaction with *Leishmania* promotes modulation of macrophage immune response to favor arginine deviation to polyamines production, leading to parasite survival. Polyamine levels are a potential modulator of growth factors, driving cells to proliferation, wound healing, oxidative stress, or regulating translation. Here, we investigate the impact of L-arginine and polyamines in the transcriptional regulation of genes involved in arginine metabolization and pro-inflammatory response to *L. amazonensis* in murine BALB/c macrophages. We found that supplementation with L-arginine is insufficient to modulate macrophage gene expression and infection. Polyamine supplementation altered nitric oxide synthase (*Nos*2) and nitric oxide (NO) production, as well as other macrophage enzymes. Putrescine supplementation increased transcript levels of polyamine metabolism-related genes Arginase 2 (*Arg*2), Spermidine synthase (*SpdS*), and Spermine synthase (*SpmS*) in both uninfected and *L. amazonensis* infected macrophages. Putrescine increased *Nos2* expression without leading to NO production, while L-arginine plus spermine led to production of NO in uninfected macrophages. Besides, L-arginine supplementation reduced the levels of *Il-1b* during infection, and L-arginine or L-arginine plus putrescine increased *Mcp1* at 24h of infection. The percentage of infected macrophages was lower after putrescine, spermidine, and spermine supplementation than L-arginine supplementation. Our data showed that the polyamines shift L-arginine-metabolism related-genes on BALB/c macrophages and affect infection by *L. amazonensis*.

## 1. Introduction

Macrophage response and polarization may determine the inflammatory process and the outcome of infection by different pathogens. Parasitic infections can modify macrophage metabolism and the activity of enzymes such as those from L-arginine and polyamines pathways. Polyamines play an important function in macrophage activation and polarization by regulating amino acid and protein synthesis, oxidative DNA damage, histone modifications, chromatin structure, and tricarboxylic acid (TCA) cycle (1–3). The polyamines putrescine, spermidine, and spermine are aliphatic polycations with low molecular weight (4). The cationic amino acid L-arginine is converted in ornithine and urea by the cytosolic enzyme arginase 1 (ARG1) and by the mitochondrial isoform arginase 2 (ARG2)(5). Ornithine is then converted by ornithine decarboxylase (ODC) into putrescine and subsequently transformed by spermidine synthase (SPDS) in spermidine, and then in spermine by spermine synthase (SPMS)(6).

The activation and polarization to pro-inflammatory M1 or anti-inflammatory M2 (pro-resolution) profiles also guide the metabolic state of macrophages (7,8). M1 macrophages are characterized by an increase in uptake of L-arginine, nitric oxide synthase 2 (NOS2) activity, and nitric oxide (NO) production in response to interferon gamma (IFN-ɣ) and lipopolysaccharide (LPS), resulting in high microbicidal activity (9,10). On the other hand, M2 macrophages are characterized by an increase in ARG1, which uses arginine to produce ornithine and subsequently polyamines, driving cell proliferation, collagen synthesis, tissue repair and wound healing (11,12). L-arginine and polyamines are valuable resources for macrophages. They are internalized via transporters from amino acid-polyamine-organocation (APC) superfamily, including cationic amino acid transporters such as CAT1/SLC7A1 and CAT2/SLCA2, and members of the solute carrier transporters (SLC’s) family (13). Interestingly, polyamines are uptaken and exported via heterodimeric transporter SLC3A2/SLC7A5 at different levels in M1/M2 polarized macrophages compared to resting cells (14–17). The SLC25A15, SLC1A5, SLC7A5, and SLC3A2 transporters regulate the uptake of L-arginine, L-glutamine, L-ornithine and polyamines in macrophages (14,15,17,18). Thus the amounts of such biomolecules could modulate the outcome of infection and alter L-arginine metabolism (19,20).

Leishmaniasis is a parasitic disease caused by *Leishmania*, endemic in the Americas, Africa, Europe, and Asia, and characterized by a large spectrum of clinical manifestations grouped into cutaneous, mucocutaneous, or visceral forms (21–23). The absence of effective treatment or vaccine reflects low public health policies commitment and is responsible for the high death rate of visceral leishmaniasis (24). *Leishmania amazonensis* is one of the causative agents of cutaneous leishmaniasis in Brazil (22,23). *Leishmania* alters between the sand fly invertebrate host from the genus *Phlebotomus* or *Lutzomyia*, and vertebrate hosts such as rodents and humans (25–28). During the bite of an infected sandfly, the promastigote form of *Leishmania* interacts with resident macrophages in the region of the injury, and is phagocytosed (29). Once inside the phagolysosome, the parasite differentiates into the amastigote form and multiplies (30).

The fate of infection by *Leishmania* is determined by the balance between macrophage activation and parasite capacity to subvert the host pro-inflammatory response by several mechanisms (31,32). One well-known mechanism is consuming L-arginine and modulating gene expression of proteins related to polyamines synthesis (20,33,34). Indeed, *Leishmania* is auxotrophic for L-arginine and other nutrients, and is able to regulate its content inside macrophages, interfering in its metabolism and consequently in microbicidal mechanisms (30,33,35–37). Besides, polyamines are essential for host-glutathione and parasite-trypanothione biosynthesis. Thus, these pathways compete for polyamines and L-arginine during infection, interfering in NO production and microbicidal activity (38,39).

Here we analyzed the influence of L-arginine and polyamines in modulating their metabolization-related genes and *L. amazonensis* infection in BALB/c-macrophages. We found that putrescine supplementation modulated *Arg*2, *Spd*S, *Spm*S and *Nos*2 transcripts without leading to NO production but reducing infectivity. On the other hand, L-arginine plus spermine increased NO production in uninfected macrophages. However, infectivity was lower after spermidine and spermine supplementation than in arginine, supporting that polyamines affect macrophage activation, interfering in the control of *Leishmania* infection, not necessarily associated with NO.

## 2. Methods

### 2.1. Ethics Statement

The experimental protocol for the animal experiments was approved by the Comissão de Ética no Uso de Animais (CEUA) from the Instituto de Biociências of the Universidade de São Paulo (approval number CEUA-IB: IB-USP 314/2018). This study was conducted in strict accordance with the recommendations in the guide and policies for the care and use of laboratory animals of São Paulo State (Lei Estadual 11.977, de 25 August 2005) and the Brazilian government (Lei Federal 11.794, de 8 October 2008).

### 2.2. Murine macrophages differentiation

Bone marrow cells were isolated from femurs and tibiae of BALB/c female mice aged 6 to 8 weeks, supplied by the Centro de Biotério da Faculdade de Medicina da Universidade de São Paulo, and maintained at the Instituto de Biociências da USP. Cells were incubated with 10% supernatant from the conditioned culture of L929 cells in RPMI 1640 medium (LGC, São Paulo, Brazil), supplemented with 10% inactivated fetal bovine serum, 50 U penicillin, 50 μg / mL streptomycin (Gibco™, USA) for 7 days at 34° C and 5% CO_2_.

### 2.3. Macrophage infection and polyamine treatment

Promastigotes of *L. amazonensis* (MHOM / BR / 1973 / M2269) were maintained in M199 medium (Gibco) supplemented with 10% heat-inactivated fetal bovine serum (Invitrogen), 5 mg/L hemine, 100 µM adenine, 100 U penicillin, 100 µg/mL streptomycin, 40 mM Hepes-NaOH, and 12 mM NaHCO3, at pH 6.85, at 25ºC for a week-long culture at low passage numbers (up to 5).

For RNA analysis, 5 × 10^6^ macrophages/well were plated in 6-well plates (SPL Lifescience, Korea) and for flow cytometry analysis, 1 × 10^6^ macrophages/well in 24-well plates. For infectivity, 2 × 10^5^ macrophages/well were plated in 8-well chamber-slides (Sigma, USA). Macrophages were maintained for 18 h at 34° C, 5% CO_2_. Then, macrophages and stationary-phase promastigote forms of *L. amazonensis* (La) were washed in 1X PBS and co-cultured at a ratio of 5 parasites per macrophage (MOI 5:1) in RPMI 1640 medium without L-arginine (R1780, Sigma-Aldrich, USA) supplemented with 2% of inactivated FBS (Invitrogen) with or without L-arginine, putrescine, spermidine, and spermine, as follows: deprived of L-arginine (arg-); supplemented with L-arginine (400 µM, arg+); supplemented with putrescine (100 µM, put+); supplemented with L-arginine plus putrescine (arg+/put+); supplemented with spermidine (100 µM, spd+); supplemented with L-arginine plus spermidine (arg+/spd+); supplemented with spermine (100 µM, spm+); supplemented with L-arginine plus spermine (arg+/spm+). These conditions were employed for uninfected (MO) and infected macrophages (MO-La).

After 4 hours of infection, the culture was washed twice with 1X PBS to remove non-phagocyted parasites and maintained with complete RPMI 1640 medium supplemented with 10% inactivated fetal bovine serum, 50 U penicillin, 50 μg / mL streptomycin at 34° C and 5% CO_2_. For the analysis of infection, the culture was maintained for 24 and 48 hours, cells in glass slides were fixed with acetone: methanol (1: 1, v: v), stained with Panoptic (Laborclin, Parana, Brazil), washed, and infectivity was analyzed by optical microscopy. The percentage of infected macrophages and the number of amastigotes per infected macrophage were calculated by randomly counting at least 500 macrophages per slide.

### 2.4. RNA extraction and Reverse Transcription

Macrophages were washed 2-times with 1x PBS. The supernatant was discarded and the pellet resuspended in 250μL of 1x PBS and 750μL of Trizol™ reagent (Invitrogen), and RNA extraction was performed following manufacturer’s instructions. The RNA was resuspended in 20μL of RNAse-free water and quantified by spectrometry (NanoDrop, Thermo Fisher Scientific). cDNA synthesis was performed using RevertAID Reverse Transcriptase kit (Thermoscientific), following the manufacturer’s instructions. Briefly, the reaction was prepared with 2μg of total RNA, 2 μL of random primer oligos (1.5 μg/ μL, ThermoScientific), 2 μL of dNTP (10mM, ThermoScientific) and water q.s.p. 26 μL and incubated at 72ºC for 5 min. Then, 8 μL of 5x Buffer, 2 μL of DTT (0.1M), 2 μL RNAse OUT and 2 μL of reverse transcriptase (200U / μL) were added, and the samples were incubated at 37ºC for 5 min, 25ºC for 10 min, 42ºC for 45min and 72ºC for 10 min. To discard possible contamination by genomic DNA, the negative controls of reverse transcription were prepared with the samples under the same conditions without reverse transcriptase. The obtained cDNAs were diluted 10 times in RNAse-free water for qPCR.

### 2.5. Relative Quantification of mRNA by RT-qPCR

The reaction was assembled with 2X SYBR Green PCR Master Mix, 200 nM of oligonucleotides and 5 μL of template cDNA (10x diluted), in a final volume of 10 μL. The reactions were performed using *StepOne* Real-Time PCR System (*Applied Biosystems*, Thermo Fisher Scientific): the first step, one cycle at 95 ° C for 10 minutes and second step, 40 cycles of 94 ° C for 30s and 60 ° C for 30s. To evaluate the qPCR efficiency, standard curves containing the target fragment cloned in pGEM T-Easy were used in 10x serial dilution from 10^8^ to 10^2^ molecules. The oligonucleotides pairs used were: *Mus musculus* - β-2-*microglobulin*-F: 5’-cactgaattcacccccactga-3 ’, β-2-*microglobulin*-R: 5’- acagatggagcgtccagaaag-3’; *Nos2*-F: 5’-agagccacagtcctctttgc-3 ’; *Nos2*-R: 5’-gctcctcttccaaggtgctt-3 ’; *Arg1*-F: 5’- agcactgaggaaagctggtc-3 ’; *Arg1*-R: 5’- cagaccgtgggttcttcaca-3 ’; *Arg2*-F: 5’- tctcctccacgggcaaattc -3 ’; *Arg2*-R: 5’-cactcctagcttcttctgccc-3 ’; *Cat1*-F: 5’-cgtaatcgccactgtgacct-3 ’; *Cat1*-R: 5’- ggctggtaccgtaagaccaa-3 ’; Slc3a2-F: 5’-gccactgagaatgcaaagacc -3 ’; *Slc3a2*-R: 5’-ttcagcacgtgatgggatgt-3 ’; *SpdS*-F: 5’-tggtggactacgcctactgt-3 ’; *SpdS*-R: 5’-tggtgcggtttttgcta-3 ’; *SpmS*-F: 5’-acactatggcagcagcaagac-3 ’; *SpmS*-R: 5’-tgtgcactgactctgtcatcc-3 ’; *Slc7a5*-F: 5’- aagggcagggattcatggtg-3’; *Slc7a5*-R: 5’-gtaggggtgtctttcagggc-3’; *Slc1a5*-F: 5’- cacttcctgtgaccttccca-3’; *Slc1a5*-R: 5’-actctagggccatggtcaatac-3’; *Slc25a15*-F: 5’- gcgaccttaaaaattgcccg-3’; *Slc25a15*-R: 5’-ctggtttctgtggaaggcga-3’; *Cat2*-F: 5’- tccaaaacgaagacaccagt-3’; *Cat2*-R: 5’-gccatgagggtgccaataga-3’; *Tnfa*-F: 5’- ccaccacgctcttctgtcta- 3’; *Tnfa*-R: 5’- agggtctgggccatagaact-3’; *Il-1b*-F: 5’- ccaagcttccttgtgcaagtg-3’; *Il-1b*-R: 5’- ctgtcaaaaggtggcatttcac-3’; *Mcp1*-F: 5’- tgatcccaatgagtaggctgg-3’; *Mcp1*-R: 5’- gcacagacctctctcttgagc-3’; *Odc1*-F: 5’- ctgccagtaacggagtccag-3’; *Odc1*-R: 5’- tcagtggcaatccgtagaacc-3’;. The fold-change was calculated by *Delta*-*Delta Ct* (ΔΔCt) method, based in β-2-microglobulin and normalized with MO/arg+ 4h group. The fold-change was presented in log2.

### 2.6. NO quantification assay

The macrophages were ungripped by incubation with 1mM EDTA in 1X PBS for 10 min at 34 ºC, and then adding RPMI plus 10% FBS and cell scraping on ice. The cells were washed by centrifugation with 1X PBS cold (500 xg, 10 min, 4ºC) and incubated with 50 μL of 5 μM DAF-FM (4-amino-5methylamino-2’,7’-dichlorofluorescein diacetate, Life Technologies, Eugene, OR, USA) diluted in 1X PBS for 30 min at 34 °C. Cells were then washed by centrifugation with 1X PBS (500 xg, 10 min, 4ºC) and resuspended in 300 μL of cold 1X PBS. Fluorescence acquisition was performed using BD Accuri cytometer (BD, Franklin Lakes, NJ, USA), and the collected data were analyzed in viable cells (PI-; FL2 detector) for 20,000 events, based on the characteristics of forward scatter (FSC) and side scatter (SSC).

### 2.7. Statistical analysis

The collected data were analyzed in GraphPad Prism software 7. The statistical analyses were performed using One-way ANOVA for mRNA and citometry analysis and Two-way ANOVA for infectivity, with a 95% confidence interval and Sidak’s test. Comparisons were established based on the groups supplemented with L-arginine uninfected and infected (MO/arg+ and MO-La/arg+, respectively). Comparisons of uninfected and infected groups were performed between similar conditions. Comparisons within a single group (uninfected or infected groups) were based on deprived x supplemented conditions for the same polyamine.

## 3. Results

### 3.1. L-arginine deprivation is not sufficient to modify gene expression of polyamine-related enzymes and transporters and NO production

To study polyamine pathway-related genes, we analyzed the effects of the precursor L-arginine on RNA abundance and NO production, by infecting BALB/c macrophage with *L. amazonensis* in conditions of L-arginine deprivation (arg-) or supplementation (arg +) for 4h. The mRNAs of genes involved in polyamine production in macrophages *Arg1* and *Arg2, Odc1, SpdS, SpmS*, and *Nos2* were analyzed by RT-qPCR and NO production was analysed by flow cytometry (Fig.1). L-arginine supplementation (arg+) was used as a reference to compare conditions of 4 and 24 h of infection. Results shown in Fig 1 indicate that under supplementation with arg transcripts levels of *Arg1, Arg2, Odc1, SpdS*, and *SpmS* were similar in uninfected (MO) and infected macrophages (MO-La; Fig. 1 A-E). However, under L-arginine deprivation, the levels of *Nos2* increased in infected macrophages at 4h and 24h compared to uninfected macrophages, but not in relation to arg+ (Fig.1F). When we compared 24 to 4 h, *Nos2* levels reduced about two times in arg+ MO and MO-La (Fig.1F). The percentage of NO producing cells (represented by DAF-FM+ cells) and median of fluorescence (MFI) of NO production didn’
st change upon infection or arginine deprivation (Fig. 1G-H). At the same time, L-arginine deprivation induced *Nos2* during infection without reflecting in the NO production.

**Figure 1:**
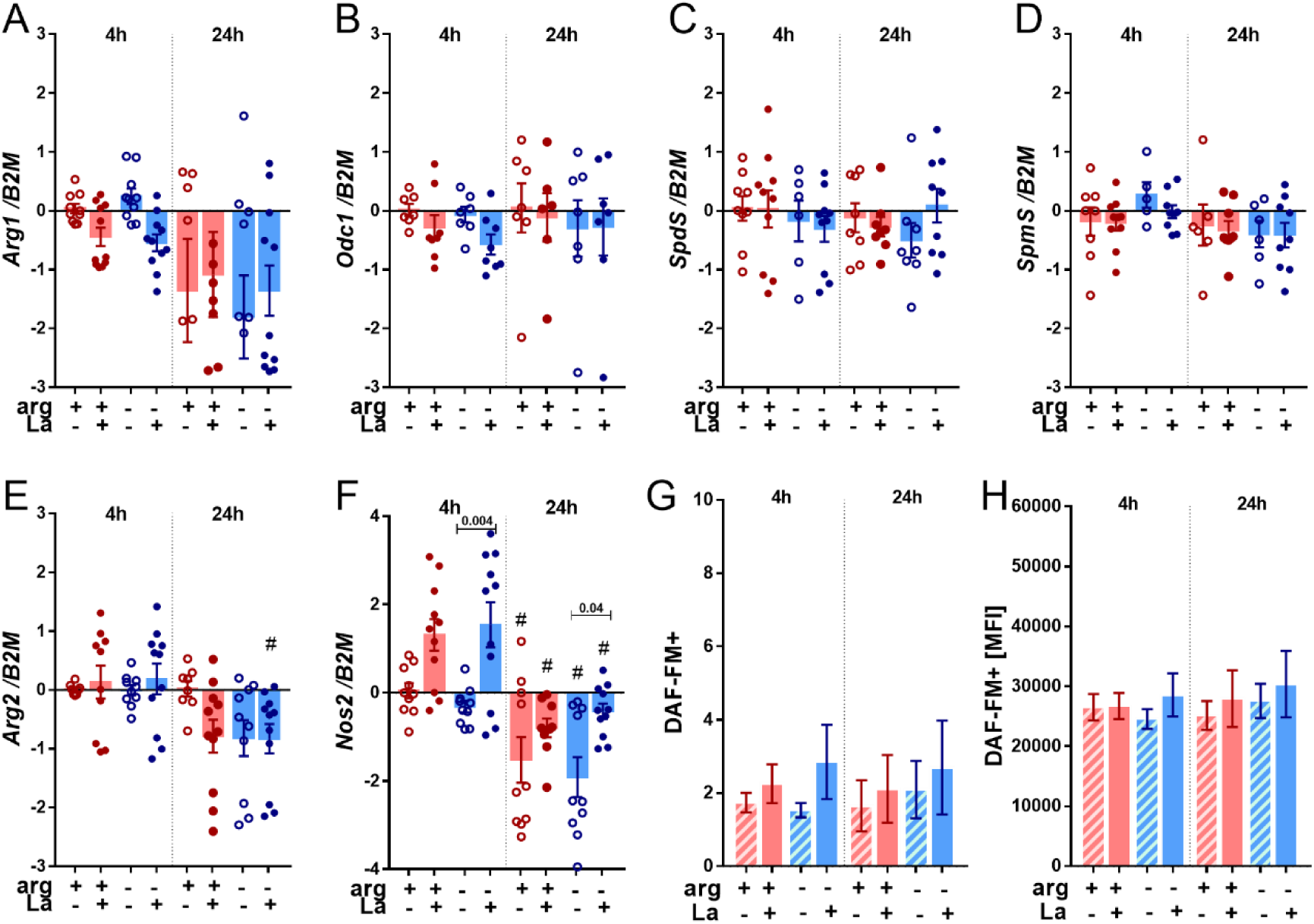
Relative expression of L-arginine metabolism related genes and NO production in BALB/c macrophages infected or not with *L. amazonensis* in conditions of L-arginine deprivation or supplementation. Macrophages were deprived of L-arginine (arg-) or supplemented with L-arginine (arg+) concomitant to *L. amazonensis* infection (MOI 5:1) for 4h and 24h. Relative quantification of *Arg1* **(A)**, *Odc1* **(B)**, *SpdS* **(C)**, *SpmS* **(D)**, *Arg2* **(E)** and *Nos2* (F) transcripts was performed by RT-qPCR. Data were normalized using *β-2-microglobulin* gene and uninfected macrophage arg+ at 4 h was used as reference for ΔΔCT relative quantification. The samples were also stained with DAF-FM to analyze the frequency of DAF-FM+ cells **(G)** and MFI **(H)** by flow cytometry. The bars represent the averages and S.E.M of the values from three independent experiments. Statistical analysis using One-Way ANOVA. **#**: p≤0.05 for the comparison between 4h vs 24h.

Considering that arginine uptake can be modulated and may alter amino acid and polyamines availability inside macrophages, we analyzed if L-arginine deprivation affected transcripts levels of the transporters of L-arginine and polyamines *Cat1 Cat2, Slc3a2*, and *Slc7a5*. As shown in Fig 2, L-arginine + and - conditions of both MO and MO-La showed reduced levels of *Cat1* and *Slc7a5* at 24 h compared to 4 h. L-arginine deprivation didn’
st modulate transcript levels for *Cat1, Cat2, Slc3a2*, and *Slc7a5* compared to arg+. These data indicate that L-arginine *per se* did not affect the expression of genes involved in L-arginine uptake and metabolism.

**Figure 2:**
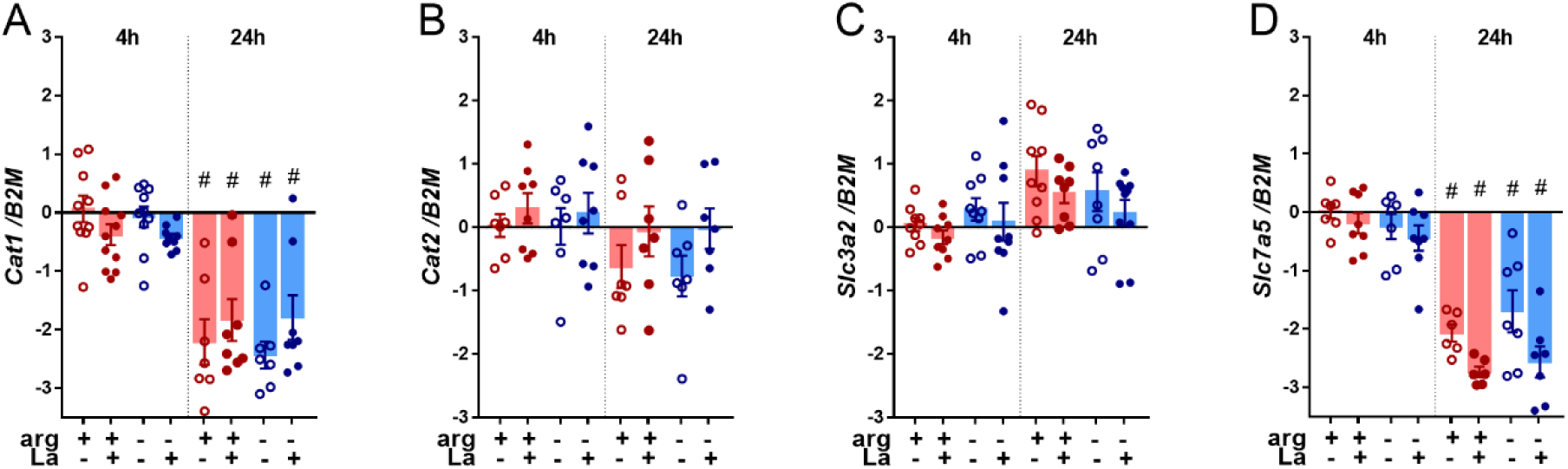
Relative expression of L-arginine and polyamines transporters in BALB/c macrophages infected or not with *L. amazonensis* in conditions of L-arginine deprivation or supplementation. Macrophages were deprived of L-arginine (arg-) or supplemented with L-arginine (arg+) concomitant to *L. amazonensis* infection (MOI 5:1) for 4h and 24h. Relative quantification of *Slc3a2* **(A)** *e Slc7a5* **(B)** transcripts was performed by RT-qPCR. Data were normalized using *β-2-microglobulin* gene and uninfected macrophage arg+ at 4 h was used as reference for ΔΔCT relative quantification. The bars represent the averages and S.E.M of the values from three independent experiments. Statistical analysis using One-Way ANOVA. **#**: p≤0.05 for the comparison between 4h vs 24h.

### 3.2. Polyamines modulate *Arg2, SpdS* and *SpmS* transcripts

To evaluate if polyamines can influence the expression of genes related to L-arginine metabolism for polyamines biosynthesis, BALB/c macrophages co-cultured or not with *L. amazonensis* for 4h were supplemented with the polyamines putrescine, spermidine and spermine in conditions of L-arginine deprivation (arg-) or supplementation (arg +).

The results shown in Fig 3 indicate that *Arg1* transcript abundances were not modulated neither by polyamines nor by infection (Fig.3A). On the other hand, L-arginine plus putrescine supplementation (arg+/put+) or putrescine only (put+) during 4h of infection led to higher levels of *Arg2* transcripts compared to only arg+ (MO-La 4h) (Fig.3B). Interestingly, the supplementation of uninfected macrophages with L-arginine plus spermidine (arg+/spd+) and spermidine only (spd+) reduced *Arg2* levels at 24h compared to arg+ (Fig.3B). A reduction of *Arg2* was also observed after supplementation of uninfected cells with L-arginine plus spermine (arg+/spm+) and spermine (spm+; Fig.3B).

**Figure 3:**
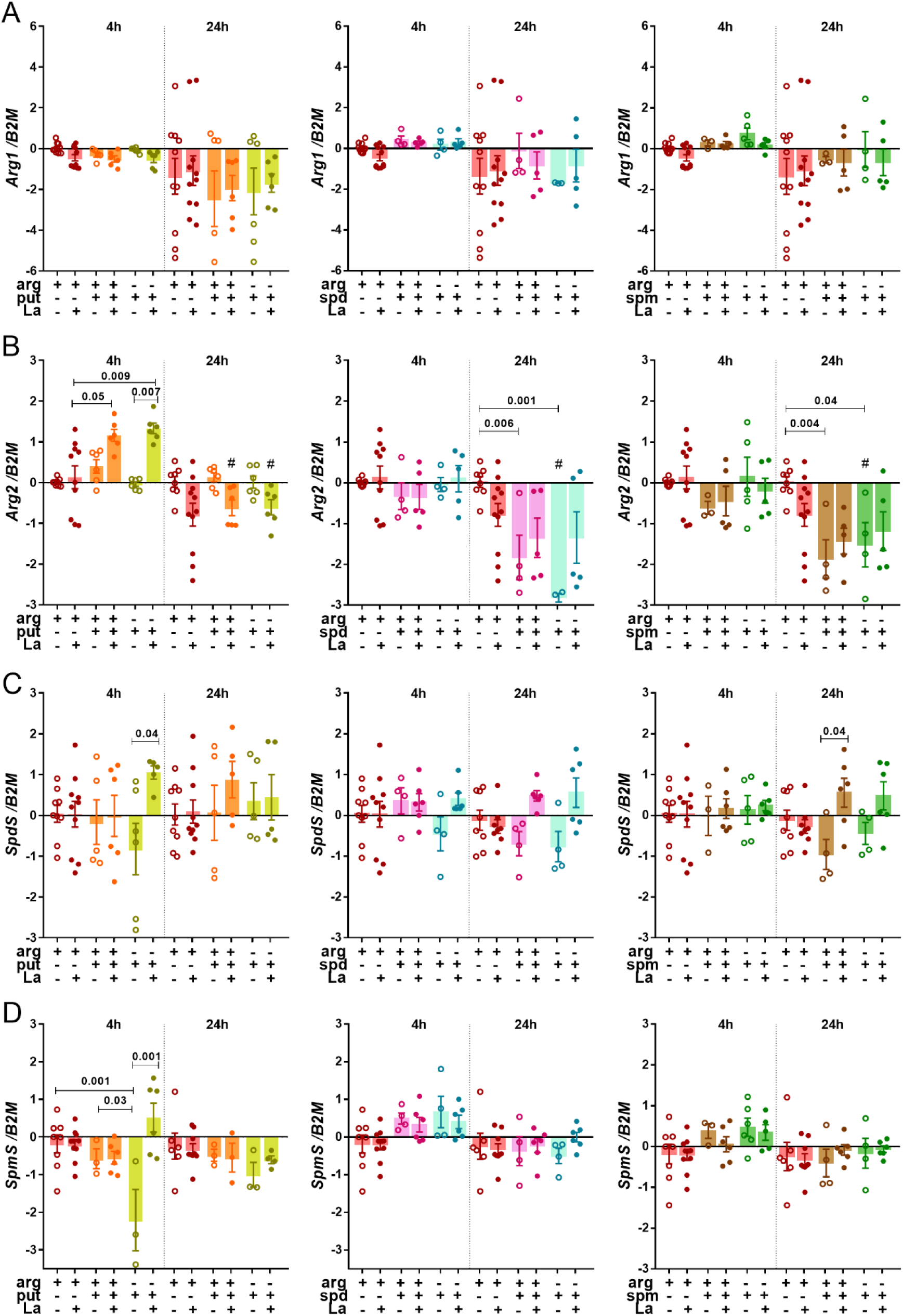
Relative expression of polyamine biosynthesis related genes in BALB/c macrophages infected or not with *L. amazonensis* with or without polyamine supplementation. Macrophages were supplemented with putrescine (put+), spermidine (spd+), spermine (spm+) with or without L-arginine (arg+) concomitant to *L. amazonensis* infection (MOI 5:1) for 4h, and after 24h in complete medium. The RNA was extracted for cDNA conversion and relative quantification of genes *Arg1* **(A)**, *Arg2* **(B)**, *SpdS* **(C)** and *SpmS* **(D)** by RT-qPCR. Data were normalized using *β-2-microglobulin* gene and the uninfected macrophage arg+ at 4 h was used as reference in ΔΔCT relative quantification. The bars represent the averages and S.E.M of the values from three independent experiments. Statistical analysis using One-Way ANOVA was indicated in the bars. **#**: p≤0.05 for the comparison between 4h vs 24h.

Comparing MO and MO-La, the supplementation with put+ and with arg+/spm increased the *SpdS* levels at 4 and at 24h, respectively (Fig.3C). In MO supplemented with put+ the levels of *SpmS* were lower at 4h compared to MO/arg+ or arg+/put+ (Fig.3D). When macrophages were supplemented with put+ during infection, we observed an increase in *SpmS* levels (Fig.3D) compared to uninfected cells.

Our data suggest that polyamines modulate the expression of *Arg2, SpdS*, and *SpmS* in both uninfected and infected macrophages.

### 3.3 Putrescine increased Nos2 expression, while spermine increased NO producing cells

We then evaluated if polyamines can influence the expression of genes related to L-arginine metabolism associated with NO production. Again, BALB/c macrophages co-cultured or not with *L. amazonensis* were supplemented with polyamines putrescine, spermidine and spermine in conditions of L-arginine deprivation (arg-) or supplementation (arg +) during 4h.

In the context of macrophage infection for 4h, supplementation with put increased more than 2-times *Nos2* levels compared to MO-La/arg+ (Fig.4A). Also, putrescine supplementation of macrophages infected for 4h increased the *Nos2* levels compared to uninfected macrophages (Fig.4A). Despite that, putrescine supplementation did not lead to increase in NO production, as stated by the similar frequencies of DAF-FM^+^ cells (Fig. 4B) and MFI values (mean of NO production per cell; Fig. 4C) in supplemented and no supplemented conditions.

**Figure 4:**
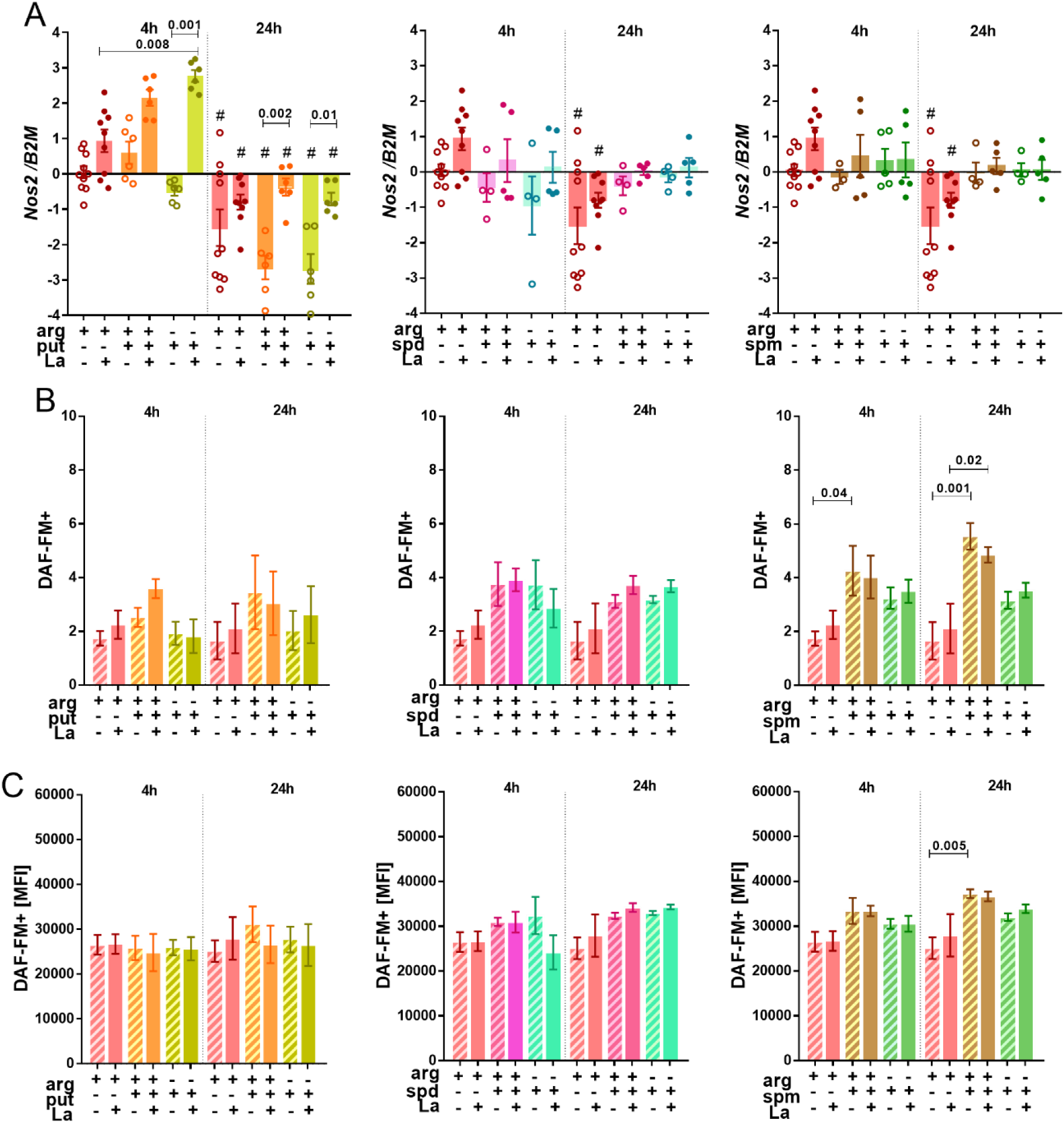
Relative expression of *Nos2* and NO production in BALB/c macrophages infected or not with *L. amazonensis* with or without polyamines supplementation. Macrophages (5×10^6^) **(A)** and (1×10^6^) **(B-C)** were supplemented with putrescine (put+), spermidine (spd+), spermine (spm+) with or without L-arginine (arg+) concomitant to *L. amazonensis* infection (MOI 5:1) for 4h, and after 24h in complete medium. The RNA was extracted for cDNA conversion and relative quantification of genes *Nos2* **(A)** by RT-qPCR. Data were normalized using *β-2-microglobulin* gene and the uninfected macrophage arg+ at 4 h was used as reference for ΔΔCT relative quantification. The samples were also stained with DAF-FM for flow cytometry analysis of DAF-FM+ cells **(B)** and MFI **(C)**. The bars represent the averages and S.E.M of the values from three independent experiments. Statistical analysis using One-Way ANOVA was indicated in the bars. **#**: p≤0.05 for the comparison between 4h vs 24h.

The supplementation with spermidine or spermine did not alter *Nos2* levels (Fig.4A). However, supplementation with arg+/spm+ increased the frequency of DAF-FM^+^ cells at 4 and 24h compared to MO/arg+ (Fig.4B). Curiously, the increase in the frequency of DAF-FM^+^ cells during 24 h macrophage infection induced by arg+/spm+ was not accompanied by changes in NO production per cell (MFI; Fig.4B-C).

Our data indicate modulation of *Nos2* expression by putrescine supplementation during infection, without a corresponding increase in NO production. On the other hand, L-arginine and spermine together can stimulate NO production in uninfected macrophages in the absence of *Nos*2 transcript modulation.

### 3.4. Polyamines affect the levels of arginine and polyamine transporters

Our next aim was to analyze if polyamines could alter the expression of genes related to L-arginine and polyamines uptake. For that, transcript levels of the transporters *Cat1, Cat2, Slc1a5, Slc3a2* and *Slc7a5* were analyzed in macrophage samples obtained under the conditions mentioned in the previous experiments.

*Cat1* levels decreased in non-infected macrophages after 24 h incubations with arg+/put+, put+, arg+/spd+, or spm compared to 4h incubations (Fig. S1A). Similarly, in infected macrophages the supplementation with arg+/put+ or put+ diminished *Cat1* levels (Fig. S1A). We also found a reduction in *Cat2* levels in uninfected macrophages supplemented with arg+/put+ in 24h compared with 4h (p≤0.006, Fig.S1B).

No differences were observed in *Slc1a5* levels between infected and uninfected cells upon supplementation (Fig. S1C). However, during macrophage infection putrescine supplementation led to increase in *Slc3a2* levels at 4h (Fig.5A). Also, the supplementation with arg/spd increased *Slc3a2* levels compared to MO-La/arg+ at 4h of infection (Fig.5A). Concerning uninfected macrophages at 24 h, arg+/spd+ or spd+ reduced the *Slc3a2* levels compared to MO/arg+ (Fig. 5A).

**Figure 5:**
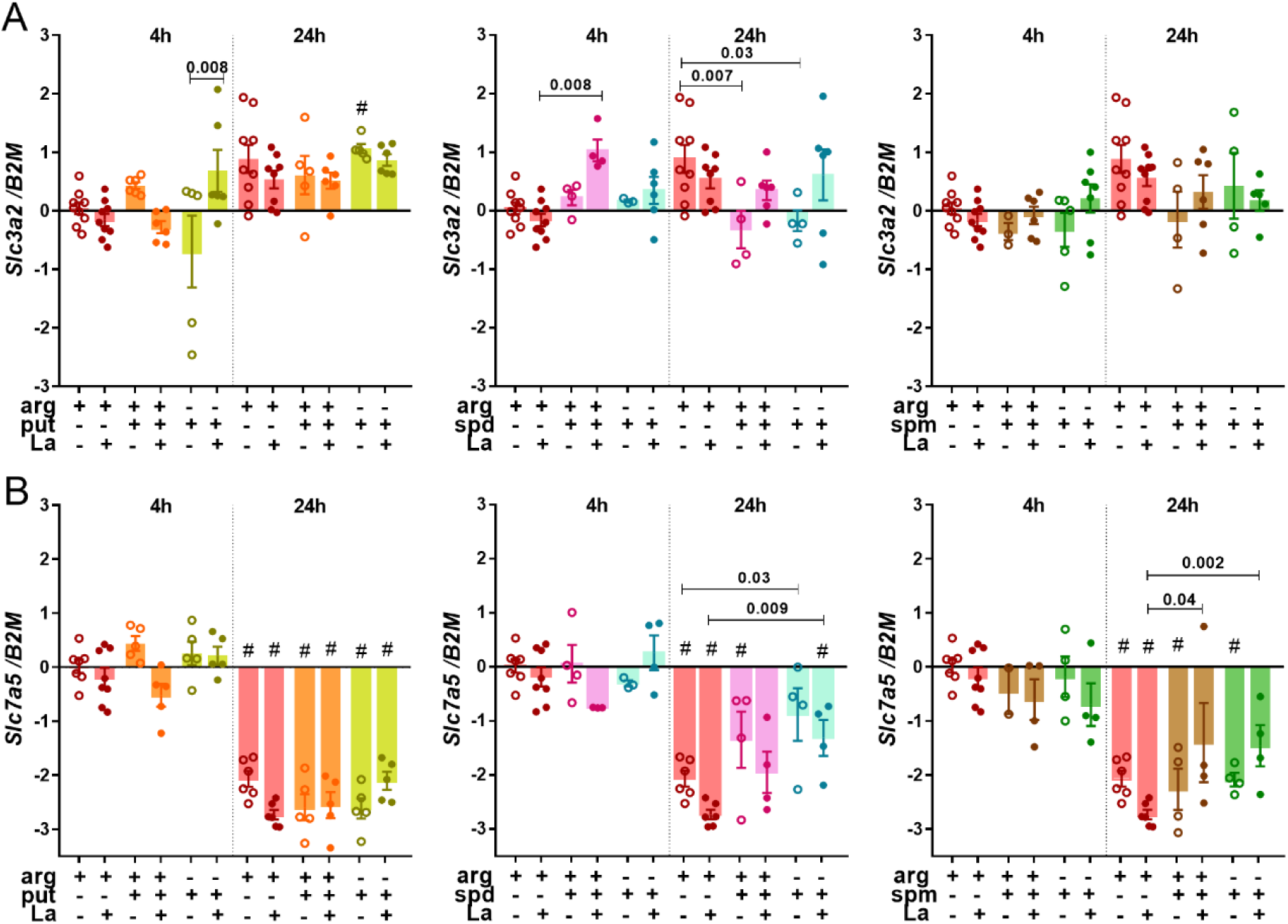
Relative expression of L-arginine and polyamines transporters in BALB/c macrophages infected or not with *L. amazonensis* with or without polyamines supplementation. Macrophages were supplemented with putrescine (put+), spermidine (spd+), spermine (spm+) with or without L-arginine (arg+) concomitant or not to *L. amazonensis* infection (MOI 5:1) for 4h, and after 24h in complete medium. RNA was extracted for cDNA conversion and quantification of *Slc3a2* **(A)** and *Slc7a5* transcripts **(B)** by RT-qPCR. Data were normalized using *β-2-microglobulin* gene and the uninfected macrophage arg+ at 4 h was used as reference for ΔΔCT relative quantification. The bars represent the averages and S.E.M of the values from three independent experiments. Statistical analysis using One-Way ANOVA was indicated in the bars. **#**: p≤0.05 for the comparison between 4h vs 24h.

We observed a reduction in the levels of *Slc7a5* at 24h compared to 4h in arg+/put+ and put+ conditions in both infected or uninfected macrophages (Fig. 5B). Indeed, most conditions of incubations with spd and spm for 24h also decreased *Slc7a5* transcript levels compared to 4h. In uninfected macrophages, the supplementation with spd+ increased the *Slc7a5* levels in comparison to MO/arg+ at 24h (Fig.5B). In infected macrophages, supplementation with spd+, spm+ or arg+/spm+ increased *Slc7a5* levels at 24h compared to arg+ (Fig.5B).

Our data indicate that supplementation with putrescine or L-arginine plus spermidine impacts the expression of *Slc3a2* in uninfected macrophages. Furthermore, spermidine supplementation alters *Slc7a5* expression in uninfected macrophages, and both spermidine or L-arginine plus spermine can modulate *Slc7a5* during macrophage infection by *L. amazonensis*.

### 3.5. Polyamines can induce *Mcp1* and *Il-1b* expression

Next, we analyzed if polyamines could alter the expression of genes of cytokines related to pro-inflammatory activation of macrophages.. Transcript levels of *Il-1b, Tnfa* and *Mcp1* were analyzed in macrophage samples obtained under the conditions mentioned in the previous experiments

In uninfected macrophages, the supplementation with arg+, arg+/put+, put+ or arg+/spd+ increased *Il-1b* levels at 24 h compared to 4h in the same conditions (Fig.6A). At 24h, infected macrophages supplemented with arg+, or arg+/put+ presented a reduction in the *Il-1b* levels compared to uninfected ones (Fig. 6A). No modifications were observed in *Tnfa* levels under arginine or polyamines supplementation during *L. amazonensis* infection, but *Tnfa* transcript levels increased in uninfected macrophages after 24h incubations with arg+/spd+ or spd+ compared to 4h (Fig.6B).

**Figure 6:**
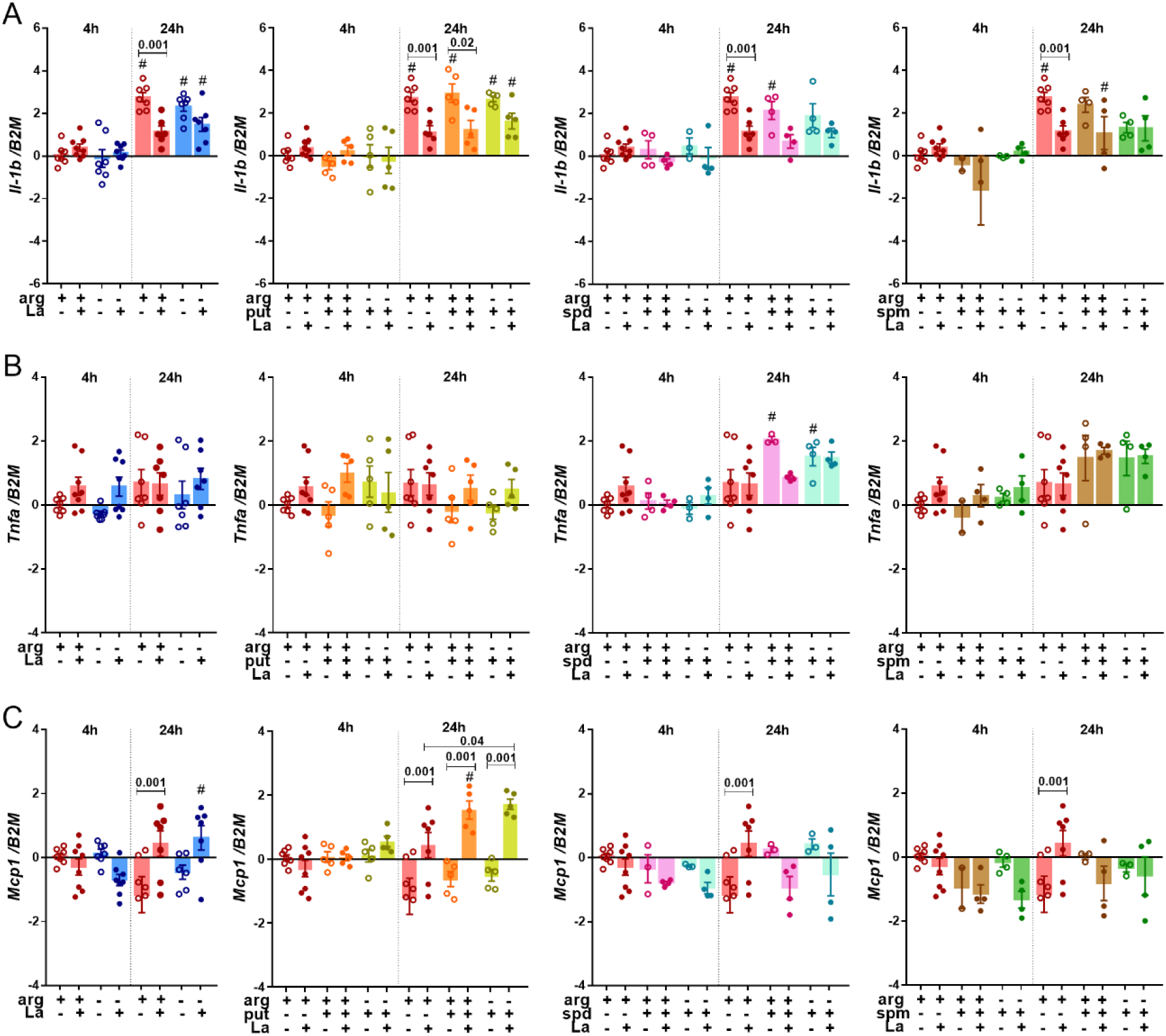
Relative expression of pro-inflammatory cytokines in BALB/c macrophages infected or not with *L. amazonensis* with or without polyamines supplementation. Macrophages were supplemented with putrescine (put+), spermidine (spd+), spermine (spm+) with or without L-arginine (arg+) concomitant or not to *L. amazonensis* infection (MOI 5:1) for 4h, and after 24h in complete medium. RNA was extracted for cDNA conversion and relative quantification of genes *Il-1b* **(A)**, *Tnfa* **(B)** e *Mcp1* **(C)** by RT-qPCR. Data were normalized using *β-2-microglobulin* gene and uninfected macrophage arg+ at 4 h was used as reference for ΔΔCT relative quantification. The bars represent the averages and S.E.M of the values from three independent experiments. Statistical analysis using One-Way ANOVA was indicated in the bars. **#**: p≤0.05 for the comparison between 4h vs 24h.

The supplementation with L-arginine increased *Mcp1* levels at 24h of infection compared to uninfected macrophages (Fig. 6C). Accordingly, *Mcp1* transcripts are significantly higher after 24h of infection compared to 4h. Macrophages infected for 24h and supplemented with arg+, arg+/put+ or put+ showed an increase in *Mcp1* levels compared to uninfected counterparts (Fig.6C).

Our data indicate that *Il-1b* and *Mcp1* transcripts can be modulated by infection in the presence of arginine and putrescine supplementation. More specifically, *Mcp1* levels increased upon infection in the presence of arginine and putrescine, while *Il-1b* levels reduced upon infection in the presence of L-arginine or L-arginine plus putrescine.

### 3.6. Polyamines and macrophage infection

We finally evaluated the impact of polyamines and arginine supplementation on the infection of macrophages by *L. amazonensis*. We determined the frequency of infected macrophages and the number of amastigotes per infected macrophage. BALB/c macrophages were co-cultured with *L. amazonensis* during 4h in the presence or not of L-arginine and polyamines, parasites were washed out, and cells were either fixed or incubated until 24h and 48h

As shown in Fig 7, the percentage of infected macrophages increased after 48 hours in the presence and in the absence of arg+ (p≤0.001) compared to 4 hours (Fig. 7A). The number of amastigotes per macrophage was similar among arg+ and arg-at all times (Fig.7B).

**Figure 7:**
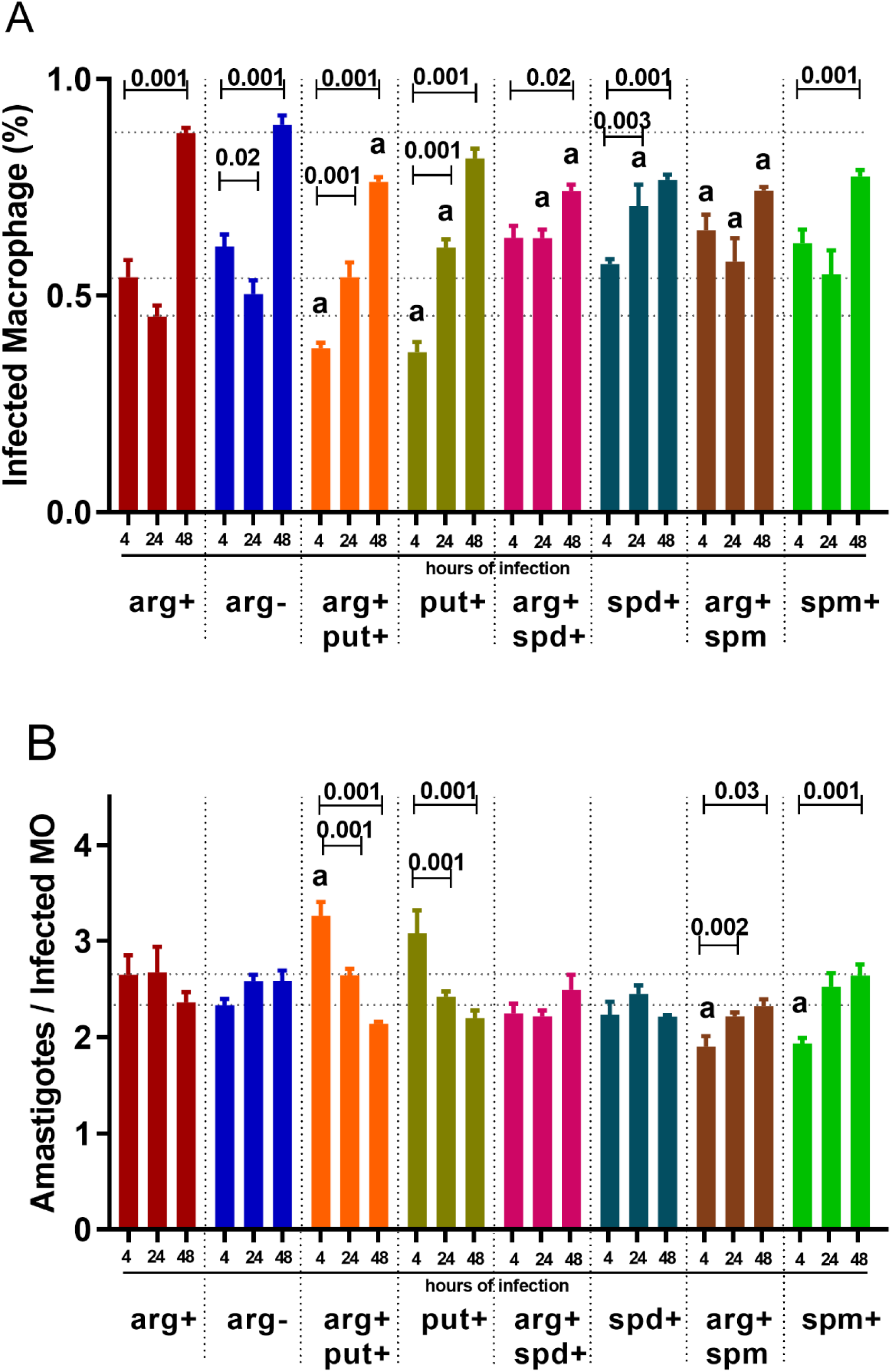
Effect of polyamine supplementation in infection of BALB/c macrophages with *L. amazonensis*. Macrophages (2×10^5^) were deprived (arg-) or supplemented (arg+) with L-arginine and/or putrescine (put+), spermidine (spd+), spermine (spm+) concomitant to *L. amazonensis* infection (MOI 5:1) for 4h, 24 and 48h. Cells were stained using Panoptic to determine the percentage of infected macrophage **(A) and**the number of amastigotes per macrophage. Each bar represents the mean ± S.E.M of the values obtained in 2 independent experiments (n = 500 macrophages). One-Way ANOVA analysis indicates less or equal values or symbols above the bars. a: p≤0,05 for the comparison between 4h vs 24h.

Unexpectedly, arg+/put+ and put+ supplementation showed a significantly lower percentage of infected macrophages at 4h compared to arg+ (Fig.7A). In arg+/put+ and put+ the percentage of infected macrophages increased at 24 and 48h compared to 4h (Fig.7A). However, upon arg+/put+ supplementation, the percentage of infected macrophages reduced at 48h compared to arg+. Curiously, the number of amastigotes per infected macrophage increased with arg+/put+ at 4h (p≤0.001) compared to arg+, but decreased after 24 and 48h.

The supplementation with arg/spd or spd increased the percentage of infected macrophages at 24h compared to arg+, but arg+/spd+ supplementation decreased the percentage of infected macrophages at 48h compared to arg+ (Fig.7B). Upon supplementation with arg and spm, the percentage of infected macrophages was higher than in arg+ at 4 and 24h, but lower at 48h. In arg+/spm+ and spm+, the number of amastigotes per infected macrophage were lower at 4h compared to arg+ at 4h (Fig.7B-C). These data indicate that polyamines interfere in a complex way in the percentage of infected macrophages and in the number of amastigotes per infected macrophage.

## 4. Discussion

Macrophages play an important role in recognizing pathogens and inducing an appropriate inflammatory activation, latter inducing T and B cell response and memory generation. The metabolization of L-arginine in macrophages can be guide to produce NO inducing cytotoxic mechanisms, or to produce polyamines. Polyamines act on the regulation of amino acid and protein synthesis, oxidative DNA damage, histone modifications, chromatin structure, and TCA cycle in macrophages (1–3), impacting on macrophage M1/M2 polarization. Studying the impact of L-arginine and polyamines metabolization in macrophages may improve our comprehension of the regulation of microbicide activity against *Leishmania*. Here, we focused on analyzing the transcript levels of genes related to L-arginine/polyamines transport and metabolism in BALB/c macrophages upon deprivation or supplementation with L-arginine, putrescine, spermidine, and spermine in uninfected conditions and during infection with *L. amazonensis*.

L-arginine availability is implicated in the outcome of *Leishmania* infection, since the competition for this amino acid by the host and parasite arginase and NOS2 affect NO production and consequently parasite killing (Fig. 8) (39–43)(42,43). Unexpectedly, we showed that deprivation or supplementation with arginine during *L. amazonensis* infection of BALB/c macrophages did not modulate the expression of *Arg1, Arg2, Odc1, SpdS*, and *SpmS*. Besides, at 4 h of infection we observed an increase and after 24h a reduction of *Nos*2 levels independent of arginine L-availability, without affecting NO production. The deprivation or supplementation with L-arginine did not alter the infection of macrophages. To estimate the effect of deprivation in L-arginine uptake we analyzed expression of arginine and polyamine transporters. Our results showed that the deprivation of this amino acid during infection did not modify the transporters *Cat1, Cat2, Slc3a2*, and *Slc7a5* levels. Interestingly, BALB/c macrophages knockout for *Cat2* present reduced transport of L-arginine during stimulation with IFN-γ plus LPS or IL-4 plus IL-10, without modifications in NOS2 or ARG1 levels (44).

**Figure 8:**
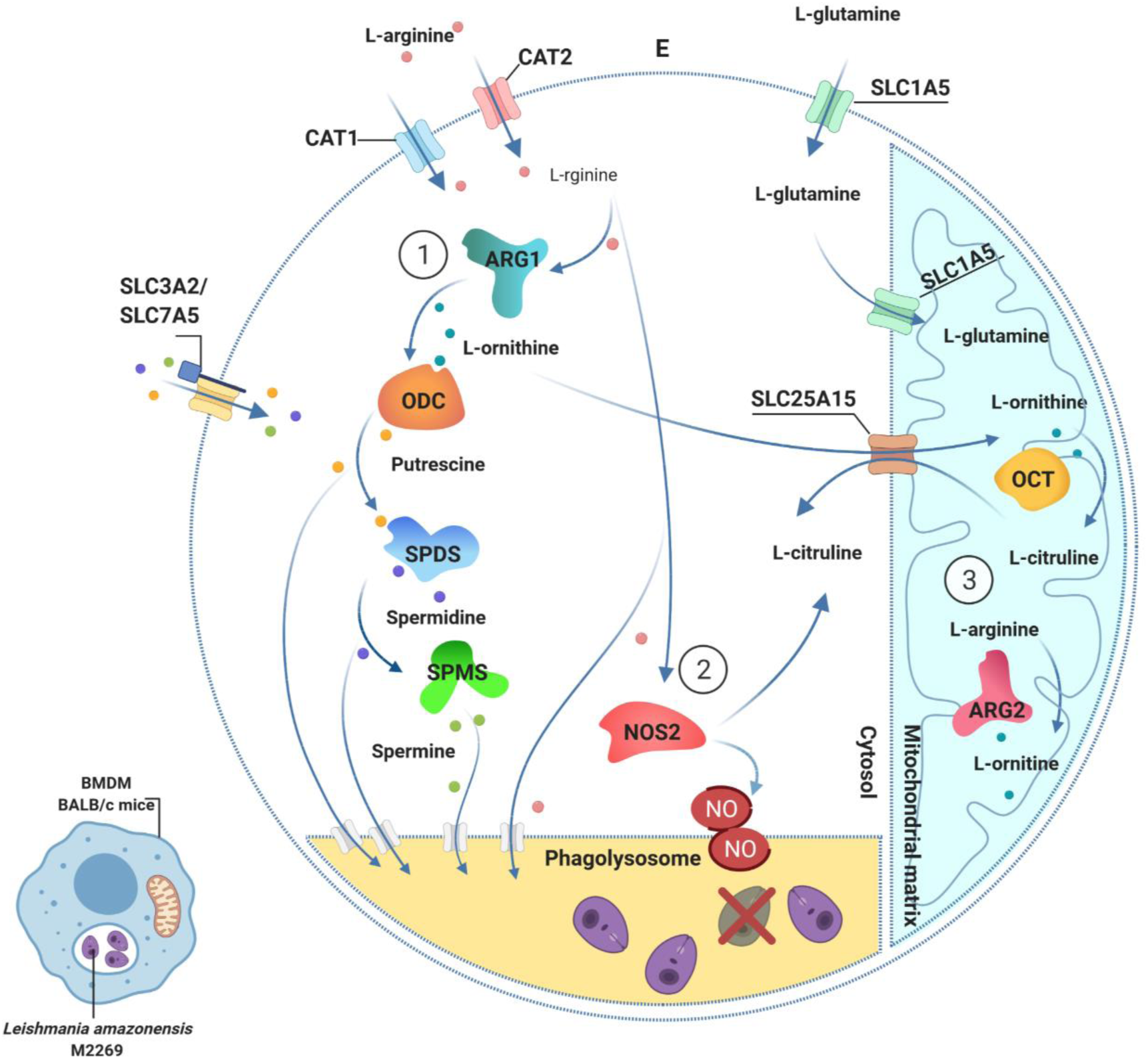
L-arginine and polyamines metabolism in BALB/c macrophage during infection with *L. amazonensis*. The L-arginine transporters CAT1 and CAT2, and the heterodimeric amino acid and polyamines transporter SLC3A2/SLC7A5 are shown in the plasmatic membrane. The amino acid and polyamines transporters can also mediate transport in phagolysosome membrane. The L-glutamine transporter SLC1A5 is shown in the mitochondrial membrane. SLC25A15 performs the L-citrulline and L-ornithine antiport between the mitochondria and cytosol. In (1) are enzymes from polyamines biosynthesis: arginase 1 (ARG1) converts L-arginine into ornithine, which is subsequently converted by ornithine decarboxylase 1 (ODC1) into putrescine. Putrescine is converted by spermidine synthase (SPDS) into spermidine; and spermidine is converted by spermine synthase (SPMS) into spermine. In (2), nitric oxide 2 (NOS2) enzyme uses L-arginine to produce nitric oxide (NO) and citrulline. In (3), mitochondrial arginase 2 (ARG2) enzyme uses L-arginine to produce ornithine. E, extracellular environment. Created with Biorender.com.

Metabolomic data reported by our group reveal increased levels of L-arginine, ornithine, putrescine, spermine and glutamine in BALB/c and C57BL/6 macrophages after 4h of infection with *L. amazonensis* (35,45). Infection of BALB/c macrophages with *L. amazonensis* knockout for arginase leads to the accumulation of L-arginine during infection, while proline, ornithine, and putrescine were diminished relative to infections with wild type parasite (35,45). These results reinforce that parasite arginase consumes L-arginine from the host, as described for amastigote forms consuming nutrients from phagolysosome, affecting the supply for ARG1 and NOS2 (46). This suggests that the internal pool of L-arginine is sufficient to support polyamines biosynthesis in the early phase of infection.

Polyamines availability can interfere in the levels of L-arginine transporters (CAT1/2) and in L-arginine metabolization by enzymes ARG1 and ARG2, and the enzymes of polyamine pathway ODC, SPDS, and SPMS (6). Also, polyamines uptake depends on the expression of polyamine transporter SLC3A2/SLC7A5 (14–17). With these facts in mind, we attempted to study the expression of macrophage metabolism-related genes upon supplementation with putrescine, spermidine and spermine, with or without L-arginine during infection. After L-arginine and putrescine or only putrescine supplementation, the levels of *Cat1, Cat2, Slc3a2, Slc7a5, Arg2, SpdS, SpmS* and *Nos2* suffered some alterations in relation to infection or time. Some of these alterations may impact in *L. amazonensis* infection outcome. The levels of *Cat1* and *Cat2* reduced in arg+/put+ and put+ supplementation during macrophage infection, implicating putrescine availability in the modulation of L-arginine transporters. The supplementation with put+ and arg+/spd+ increased *Slc3a2* at 4h and reduced at 24h in uninfected macrophages. Also, spd+ increased *Slc7a5* at 24h in uninfected macrophages. During *L. amazonensis* infection, spermidine or L-arginine plus spermine can modulate *Slc7a5*, suggesting that polyamines can influence *Slc3a2/Slc7a5* transcription.

We have already reported that higher levels of citrulline in BALB/c and C57BL/6 macrophages after 4h of infection with *L. amazonensis* did not correlate with NO production, once the NO production was observed only in C57BL/6 macrophages (35,45,47). We now showed that the supplementation with putrescine increased *Nos2* levels, but did not reflect in increased NO production. NO producing cells were observed in L-arginine plus spermine supplementation in both uninfected and *L. amazonensis* infected conditions, without *Nos2* modulation. Curiously, another group reported that spermidine reduced the expression of *Nos*2 in LPS-stimulated macrophages (48). It was previously shown that BALB/c macrophages knockout for *NOS2* did not differ in the expression of *Cat1* and *Cat2*, and also transport L-arginine during stimulation with IFN-γ plus LPS or IL-4 plus IL-10 (44). Our datacan support the idea that polyamines can exert a major influence in NO production during infection.

In BALB/c and C57BL/6 macrophages after 4h of infection with *L. amazonensis* we previously observed increase in ornithine, putrescine, spermine, proline and glutamine levels (35,45). Through the urea cycle, ornithine can be metabolized to produce proline and glutamine (49–51). In contrast to our expectation, we did not find modulation in the levels of *Arg1* upon polyamine supplementation. On the other hand, *Arg2* abundance increased in the presence of L-arginine plus putrescine or only putrescine in infected-macrophages after 4h. Despite the lack of polyamines biosynthetic pathways in mitochondria, we can speculate that increased level of *Arg*2 induced by putrescine availability can guide the use of L-arginine by ARG2 increasing ornithine, which could be converted in citrulline inside mitochondria or polyamines in cytoplasm. The increased levels of ornithine, proline and glutamine in *L. amazonensis* infected BALB/c macrophages can support glutamate production, interfering in the metabolic state of macrophages during activation (31). In M2 macrophages, the activation of the oxidative phosphorylation system (OXPHOS) is associated with releasing polyamines, allowing cell proliferation and wound healing (49,52,53). On the other hand, glycolytic metabolism can be induced in macrophages by the availability of O_2_, glucose, glutamine, L-arginine, and polyamines to generate ATP, characteristic of aerobic metabolism in pro-inflammatory M1 macrophages. The polyamines contribute to glutamate production and subsequent α-ketoglutarate production, broking the tricarboxylic acid cycle (TCA) in the Warburg effect and affecting ATP production and the redox state of cells (49,52,53).

Spermine and spermidine play a role in protecting cells from reactive oxygen species (ROS). Spermidine is know to reduce the expression of ROS in LPS-stimulated macrophages (48). Polyamines, specially spermine, can indirectly mediate Ca^2+^ transport or function on mitochondrial respiration, stimulating succinate dehydrogenase activity, coupling an increased mitochondrial reactive oxygen species (mtROS) production (54–56). Glutathione is composed by glycine, cysteine and glutamic acid and acts as a ROS scavenger in thiol-disulfide reactions to control the protein functions (57). *Leishmania* can use glutathione and spermidine to produce glutathionylspermidine, and subsequently trypanothione, an essential molecule to protect the parasite against the mammalian host defense (58,59). Here, we found that uninfected macrophages supplemented with L-arginine plus spermine displayed an increase in NO production. Despite that, the infection subverts this effect suggesting that *L. amazonensis* blocks the macrophage capacity to respond to the variations in polyamine levels, deviating the macrophage activity to ensure its survival. Spermine induces superoxide dismutase synthesis and can prevent oxidative damage (60,61). The dysregulation of antioxidant activity leads to ROS accumulation and affects mitochondrial integrity (62). Spermine also negatively regulates macrophage activation, which can be linked to polyamines catabolism mediated by acetylation via N1-spermidine/spermine acetyltransferase (SSAT) (63,64).

In uninfected macrophages, putrescine supplementation reduced expression of *SpmS* at 4h compared to L-arginine supplementation. Expression was restored upon macrophage infection, being similar to the condition of L-arginine supplementation. This observation suggests that *L. amazonensis* blocks/interferes in the macrophage capacity to respond to variations in polyamine levels, possibly to ensure its survival inside the host. On the other hand, the accumulation of polyamines can cross-regulate metabolic-related genes during infection. Also, it was shown that *SpmS* knockout cause accumulation of spermidine and increase of aldehyde and hydrogen peroxide (H_2_O_2_), leading to lysosomal dysfunction and oxidative stress (61). Spermine was also shown to inhibit translation of NOS2 mRNAs in macrophages, reducing NO production (55). IL-4 stimulation induces ODC, increasing putrescine production in murine macrophages, and the inhibition of ODC with difluomethylornithine (DFMO) reduces putrescine content but not spermidine and spermine (65,66). In this context, it’s interesting to cite the function of spermidine in the regulation of translation by mediating spermidine-hypusination of lysine residue in eukatiotic initiation factor (eIF5A) (38,39). Also, macrophages stimulated with IL-4 display an increase in putrescine levels and hypusination of eIF5A (38,39).

Regarding the expression of genes related to macrophage polarization, putrescine supplementation during infection increased the expression of *Nos2* and *Mcp1* independently of L-arginine supplementation. Previous studies showed that the putrescine and spermine increase MCP-1 and TNF-α in mixed glial culture (67). Spermidine reduces the secretion of TNF-α and IL-1β in LPS-stimulated RAW 264.7 macrophages (48), and MCP-1 secretion in THP-1-macrophages treated with IFN-γ (68).

Our results suggest that polyamines can act as protagonists in activating of inflammatory response to *Leishmania* infection, not only by releasing L-arginine and inducing NO production, but also by acting in the host-glutathione and redox balance, improving the microbicide activity of macrophage. Putrescine, spermidine and spermine supplementation reduced the percentage of infected macrophages, corroborating the idea that polyamines favor macrophage activation and microbicide capacity, to the detriment of parasite survival.

## 6. Author Contributions

Conceptualization, J.M.Z. and S.M.M.; methodology and experiments design, J.M.Z., S.M.A., and C.A.B.; designed the figure; J.M.Z., S.M.A. and S.M.M.; writing—original draft preparation, J.M.Z., B.S.C and S.M.M; All authors have read and agreed to the published version of the manuscript.

## 8. Acknowledgments

We thank to Professor Jean Pierre Schatzmann Peron and Lab members for welcoming, encouraging and giving support to finish this work: Carolina Manganeli Polonio, Nagela Ghabdan Zanluqui, Marília Garcia, Lilian Gomes de Oliveira, Laura Caroline de Faria, Tiago Francisco da Silva, Yan Souza Angelo and Igor Santiago Carvalho.

