## Supplementary material for "Polyamines shift expression of macrophage L-arginine metabolism related-genes during *Leishmania amazonensis* infection"

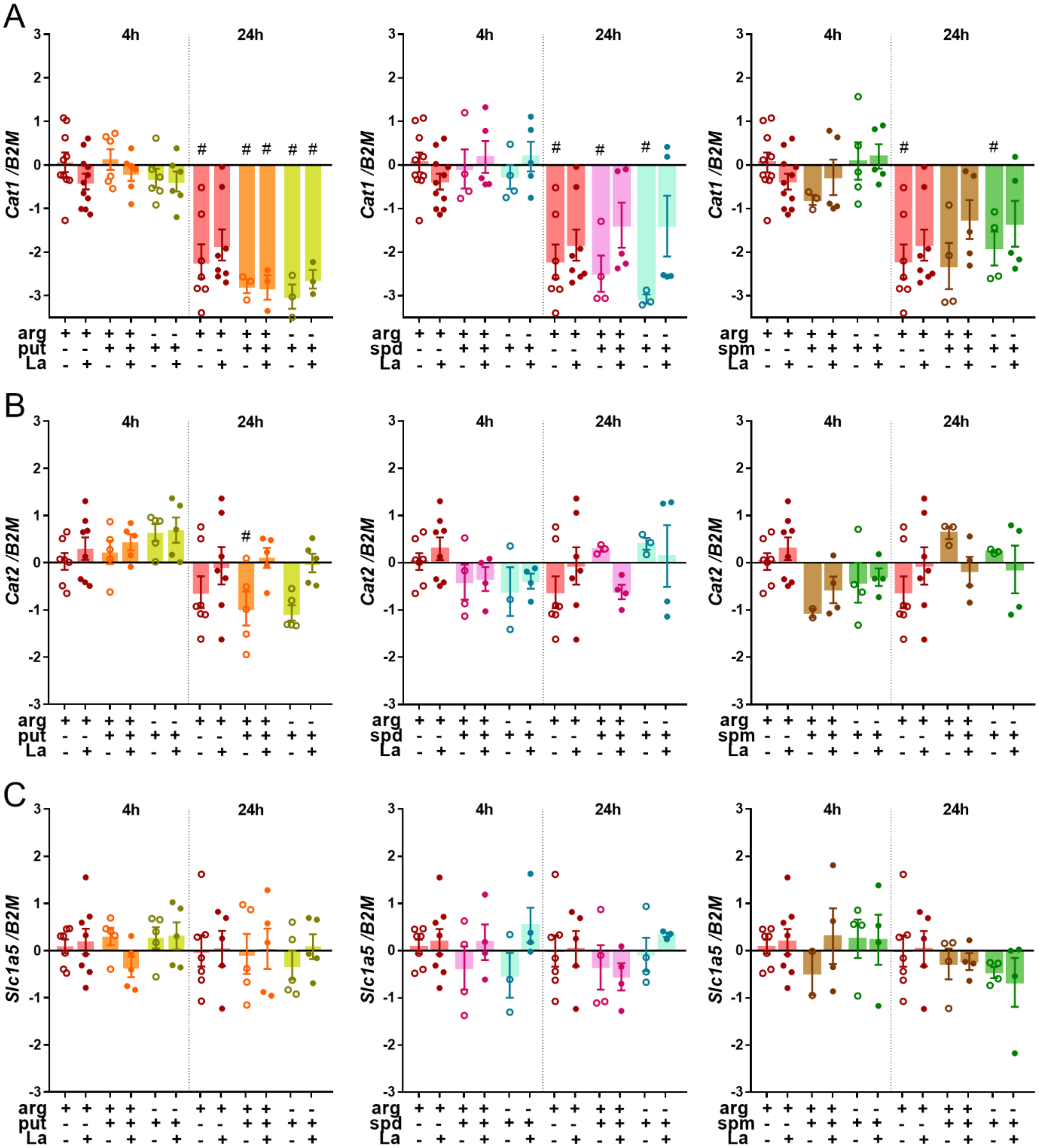

**Figure S1: Gene expression of cationic and L-glutamine transporters in uninfected and infected BALB/c macrophages**. The macrophages ($5x10^6$) were supplemented with L-arginine (arg+) and/or putrescine (put+), spermidine (spd+), spermine (spm+), simultaneously to *L. amazonensis* infection, maintained in the MOI proportion of 5:1 for 4h and, after, to more 24h in complete medium. After 4 and 24h, the RNA was extracted for cDNA conversion and relative quantification of genes *Cat1* **(A)**, *Cat2* **(B)** and *Slc1a5* **(C)** by RT-qPCR. The data were normalized using the *β-2-microglobulin* gene. The uninfected macrophages supplemented with arg+ at 4h was used as control in ΔΔcT calcules. The bars represent the averages and S.E.M of the values. One-Way ANOVA analysis indicates less or equal values or symbols above the bars. #: p≤0,05 for comparing 4h vs. 24h.

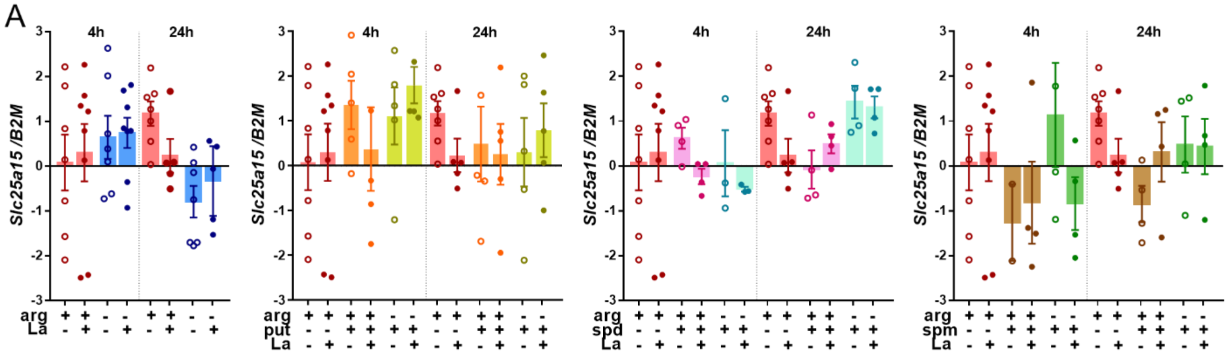


**Figure S2: L-citrulline and L-ornithine antiporter gene expression in uninfected and infected BALB/c macrophages**. The macrophages ($5x10^6$) were supplemented with L-arginine (arg+) and/or putrescine (put+), spermidine (spd+), spermine (spm+), simultaneously to *L. amazonensis* infection, maintained in the MOI proportion of 5:1 for 4h and, after, to more 24h in complete medium. After 4 and 24h, the RNA was extracted for cDNA conversion and relative quantification of the gene *Slc25a15* **(A)** by RT-qPCR. The data were normalized using *β-2-microglobulin* gene. The bars represent the averages and S.E.M of the values. One-Way ANOVA analysis indicates less or equal values or symbols above the bars. #: p≤0,05 for comparing 4h vs. 24h.
